# A fast multi-locus random-SNP-effect EMMA for genome-wide association studies

**DOI:** 10.1101/077404

**Authors:** Yang-Jun Wen, Hanwen Zhang, Jin Zhang, Jian-Ying Feng, Bo Huang, Jim M. Dunwell, Yuan-Ming Zhang, Rongling Wu

**Affiliations:** State Key Laboratory of Crop Genetics and Germplasm Enhancement, Nanjing Agricultural University, Nanjing 210095, China; Faculty of Applied Science, The University of British Columbia, Vancouver V6T 1Z4, Canada; School of Agriculture, Policy and Development at University of Reading, Reading RG6 6AR, United Kingdom; Statistical Genomics Lab, College of Plant Science and Technology, Huazhong Agricultural University, Wuhan 430070, China; Center for Statistical Genetics, The Pennsylvania State University, Hershey, PA 17033-0850, USA

**Keywords:** genome-wide association study, mixed linear model, multi-locus mapping, random effect

## Abstract

Although the mixed linear model (MLM) such as efficient mixed model association (EMMA), has been widely used in genome-wide association studies (GWAS), relatively little is known about fast and efficient algorithms to implement multi-locus GWAS. To address this issue, we report a fast multi-locus random-SNP-effect EMMA (FASTmrEMMA). In this method, a new matrix transformation was constructed to obtain a new genetic model that includes only quantitative trait nucleotide (QTN) variation and normal residual error; letting the number of nonzero eigenvalues be one and fixing the polygenic-to-residual variance ratio was used to increase computing speed. All the putative QTNs with the ≤0.005 P-values in the first step of the new method were included in one multi-locus model for true QTN detection. Owing to the multi-locus feature, the Bonferroni correction is replaced by a less stringent selection criterion. Results from analyses of both simulated and real data showed that FASTmrEMMA is more powerful in QTN detection, model fit and robustness, has less bias in QTN effect estimation, and requires less running time than the current single- and multi-locus methodologies for GWAS, such as E-BAYES, SUPER, EMMA, CMLM and ECMLM. Therefore, FASTmrEMMA provides an alternative for multi-locus GWAS.

---

Genome-wide association studies (GWAS) have been widely used in the genetic dissection of quantitative traits in human, animal and plant genetics, especially in combination with the output of genomic sequencing technologies. The most popular method for GWAS is the mixed linear model (MLM) method [1–2], because of its demonstrated effectiveness in correcting the inflation from many small genetic effects (polygenic background) and in controlling the bias of population stratification [3–7]. Since the MLM of Yu *et al.* [2] was published, many MLM-based methods have been proposed. However, most of them comprise a one-dimensional genome scan by testing one marker at a time, which is involved in multiple test correction for the threshold value of significance test. The widely-used Bonferroni correction is often too conservative to detect many important loci for quantitative traits.

Most quantitative traits are controlled by a few genes with large effects and numerous polygenes with minor effects. However, the current one-dimensional genome scan approaches for GWAS do not match the true genetic model for these traits. To overcome this issue, multi-locus methodologies have been recommended, for example, Bayesian LASSO [8], adaptive mixed LASSO [9], penalized Logistic regression [10–11], Elastic-Net [12], empirical Bayes [13] and empirical Bayes LASSO [14]. If the number of markers is several times larger than the sample size, all marker effects can be included in one single model and estimated in an unbiased way. If the number of markers is much many times larger than the sample size, however, these shrinkage approaches will fail. In this situation, we should consider how to reduce the number of marker effects in the multi-locus genetic model. For example, Zhou *et al.* [15] developed a Bayesian sparse linear mixed model and Moser *et al.* [16] proposed a Bayesian mixture model. Under these models, two to four common components in the mixture distribution were considered and only a few variance components were estimated. Although about 500 effects in the genetic model are finally considered after several rounds of Gibbs sampling, the computing time becomes a major concern for these Bayesian approaches. Recently, Segura *et al.* [17] and Wang *et al.* [7] have proposed multi-locus MLM approaches. However, further refinement for fast algorithm is needed.

In the MLM methods of Zhang *et al.* [1], the QTN effect is viewed as random so that three component variances (QTN, polygenic and residual variances) need to be estimated. If the number of markers is large, this calculation takes a long time. To reduce computing time and increase power in QTN detection, a compressed MLM (CMLM) with a P3D algorithm [18] and an enriched CMLM (ECMLM) [19] have been proposed. On the other hand, Kang *et al.* [3]proposed an efficient mixed model association (EMMA), and other authors suggested alternatives, such as EMMA eXpedited (EMMAX) [20], FaST-LMM [21], FaST-LMM-Select [22], genome-wide EMMA [4] and GRAMMA-Gamma [23]. Recently, SUPER [24] has been developed based on FaST-LMM. Among the above fast methods, the SNP effect was viewed as fixed. It should be noted that Goddard *et al.* [25] treated marker effects as random, due to several advantages of the random model approach over the fixed model treatment [7,25–27]. For example, the random model approach will shrink the estimated SNP effects toward zero. However, Goddard *et al.* [25] did not provide an efficient computational algorithm to estimate marker effects. Although the random-SNP-effect MLM [7,27] can solve this issue, a new and efficient algorithm for fast calculation is also required.

In this paper, first, we treated the SNP effect as random. Although three component variances need to be estimated, a fast and new matrix transformation in the new method was constructed in order to quickly scan each marker on the genome. Then, all the putative QTN with a ≤0.005 P-value in the first step of the new method were placed into one multi-locus genetic model and these QTN effects were estimated by EM empirical Bayes (EMEB) [28] for true QTN identification. This new method, called fast multi-locus random-SNP-effect EMMA (FASTmrEMMA), was validated by analysis of real data from *Arabidopsis* [29] and by a series of simulation studies, as compared with the other methods: E-BAYES (multi-locus model) [30], SUPER, EMMA, CMLM and ECMLM (single-locus model).

The new methods have been implemented in R. The R software FASTmrEMMA is available.

## RESULTS

### Fast multi-locus random-SNP-effect EMMA (FASTmrEMMA)

#### Estimation of the QTN variance

FASTmrEMMA is a new algorithm that can approximate the estimation of QTN variance (see Materials and Methods). Thus, we need to know whether this approximation has a significant effect on the estimate of QTN variance. To answer this question, four flowering time traits in *Arabidopsis* [29] were analyzed by FASTmrEMMA and an exact method implemented by PROC MIXED in SAS. The estimates for QTN variance are listed in **Figure 1** and **Table S1**. As a result, the relative error between the two methods ranged from 0.0 to 24.09%, and the average was 1.60%, indicating no effect on the QTN variance estimate using FASTmrEMMA under the conditions of this simulation.

**Figure 1.**
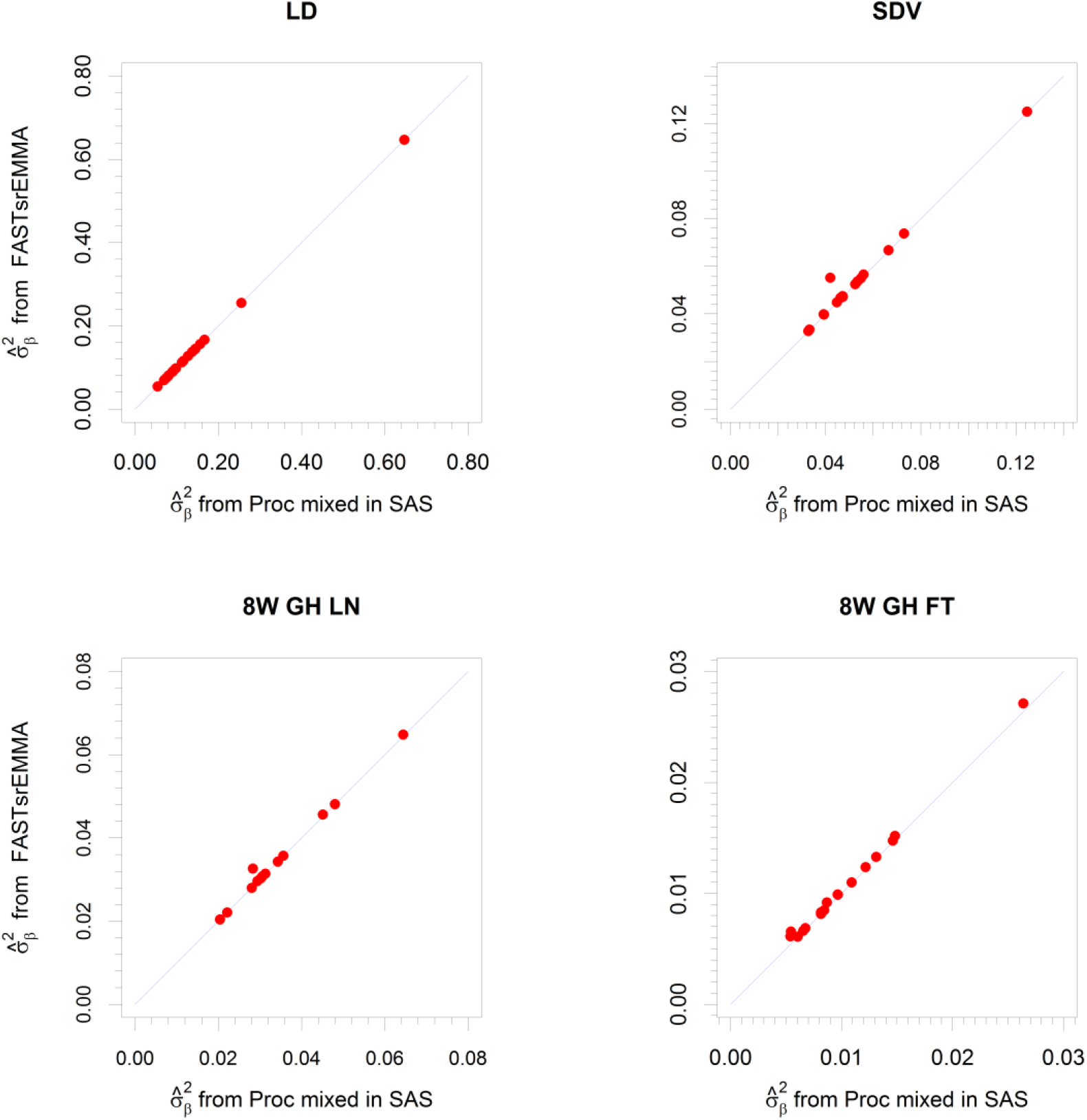
Comparison of the QTN-variance estimates between fast single-locus random-SNP-effect EMMA (FASTsrEMMA) and one exact algorithm implemented by PROC MIXED in SAS. LD: days to flowering under long days; SDV: days to flowering under short days with vernalization; 8W GH LN: leaf number at flowering with 8 wks vernalization, greenhouse; and 8W GH FT: days to flowering, 8 wks vernalization, greenhouse.

To confirm the effectiveness of FASTmrEMMA, three Monte Carlo simulation experiments were carried out and the simulation procedures were same as those in Wang *et al.* [7]. Each sample in these simulation experiments was analyzed by six methods. In the six methods, FASTmrEMMA is also a new multi-locus algorithm within the framework of MLM, E-BAYES [30] is an existing multi-locus approach under the framework of Bayesian statistics, and SUPER, EMMA, ECMLM and CMLM are the existing single-locus GWAS methods.

#### Statistical power for QTN detection

In the above three simulation experiments, the power for each QTN was defined as the proportion of samples where the QTN was detected (the P-value is smaller than the designated threshold). When only six QTNs were simulated in the first experiment, the power in the detection of each QTN was higher for FASTmrEMMA than for the others (**Figure 2a**; **Table S2**). When a polygenic background (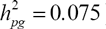) was added to the first experiment, a similar trend was observed (**Figure 2b**; **Table S2**). On one occasion (QTN number 1), E-BAYES method was slightly more powerful than FASTmrEMMA method. When the polygenic background was changed into an epistatic background (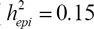), the results were also similar to those in the first experiment (**Figure 2c**; **Table S2**). These results demonstrate the highest power of FASTmrEMMA across all the approaches under various genetic backgrounds, although the other methods are also robust under these backgrounds.

**Figure 2.**
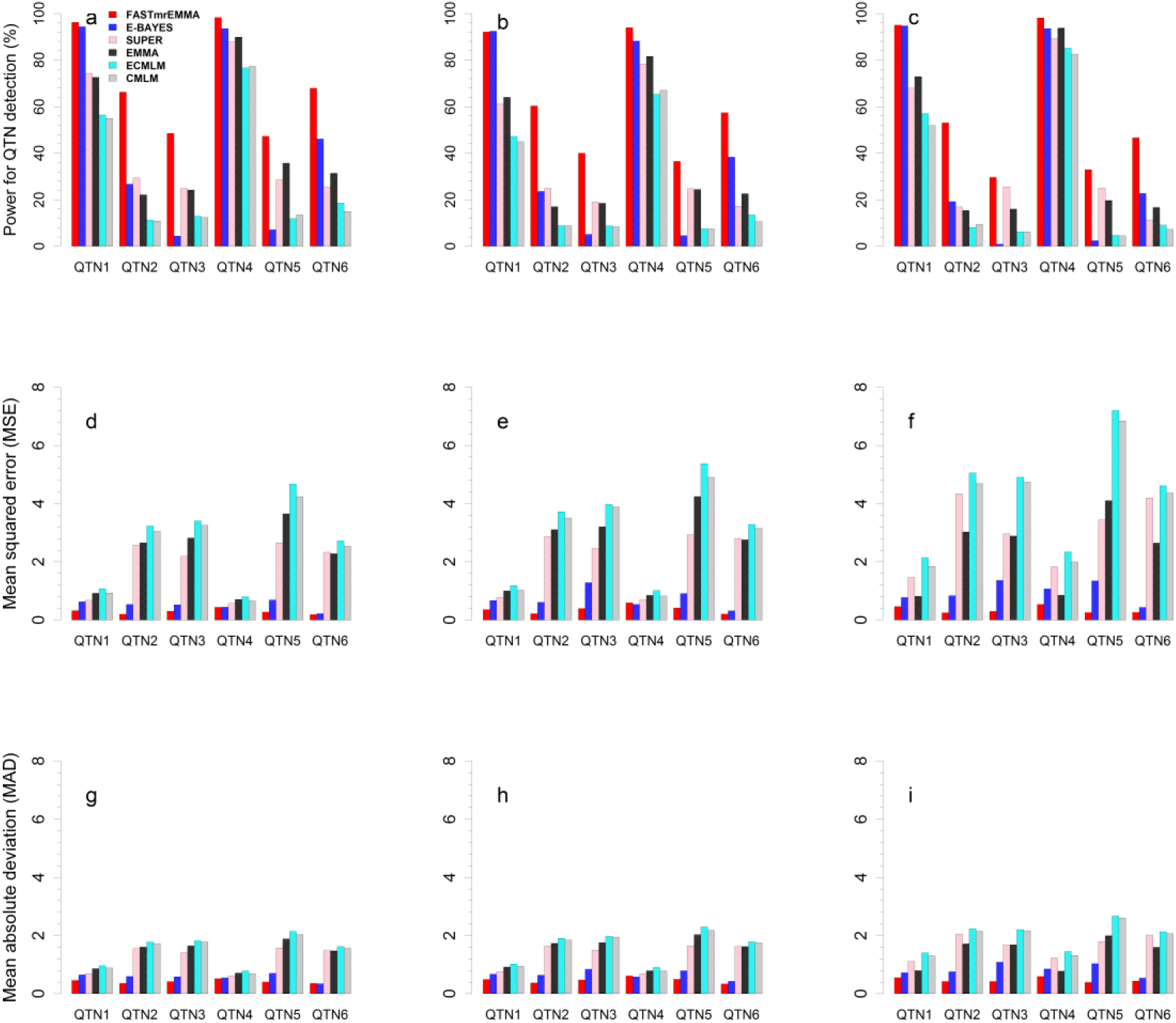
Comparison of fast multi-locus random-SNP-effect EMMA (FASTmrEMMA) with the single- and multi-locus approaches under various genetic backgrounds. The single-locus model approaches include SUPER, EMMA, CMLM and ECMLM, and the multi-locus approach has E-BAYES. The powers are presented in a-c, mean squared errors are showed in d-f, and mean absolute deviations are listed in g-i. Six QTN (a, d and g), six QTN plus polygenes (b, e and h), and six QTN plus three epistasis (c, f and i) were simulated, respectively, in the second to fourth simulation experiments.

#### Accuracy for estimated QTN effects

We used the average, MSE and MAD to measure the accuracy of an estimated QTN effect. We evaluated the accuracies for the estimates of all the six simulated QTNs across all the six methods. As a result, the estimate of each QTN effect from FASTmrEMMA was much closer to the true value than the estimates obtained from the other methods. On these occasions (QTN numbers 1 and 4), the averages from E-BAYES were closer to the true value than those from FASTmrEMMA in three simulation experiments (**Table S2**). The MSE and MAD for each QTN effect were significantly less from FASTmrEMMA than from the others with an exception for QTN number 4 where E-BAYES has slightly less MSE and MAD than FASTmrEMMA in the third simulation experiment (**Figures 2d-2i**; **Table S2**). These results indicate that a higher accuracy for the estimate of QTN effect can be achieved using FASTmrEMMA than using the other methods.

#### False Positive Rate and ROC curve

All the false QTN, detected by the six methods, in three simulation experiments were used to calculate the empirical false positive rates of the six methods. These results are listed in **Table S3**. In these three simulation experiments, the empirical false positive rates of the six methods were between 0.096 and 5.181 (× 1E-4), and had the same order of magnitude. EMMA has the lowest false positive rate followed by ECMLM, CMLM and FASTmrEMMA methods, and SUPER has the maximum false positive rate followed by E-BAYES method.

A receiver operating characteristic (ROC) curve is a plot of the statistical power against the controlled type I error. This curve is frequently used to compare different methods for their efficiencies in the detection of significant effects; the higher the curve, the better is the method. When eleven probability levels for significance, between 1e-8 to 1e-3, were inserted, the corresponding powers were calculated in the second simulation experiment. The results are shown in **Figure 3**. Among the six approaches, clearly, FASTmrEMMA method is the best one and the next one is E-BAYES.

**Figure 3.**
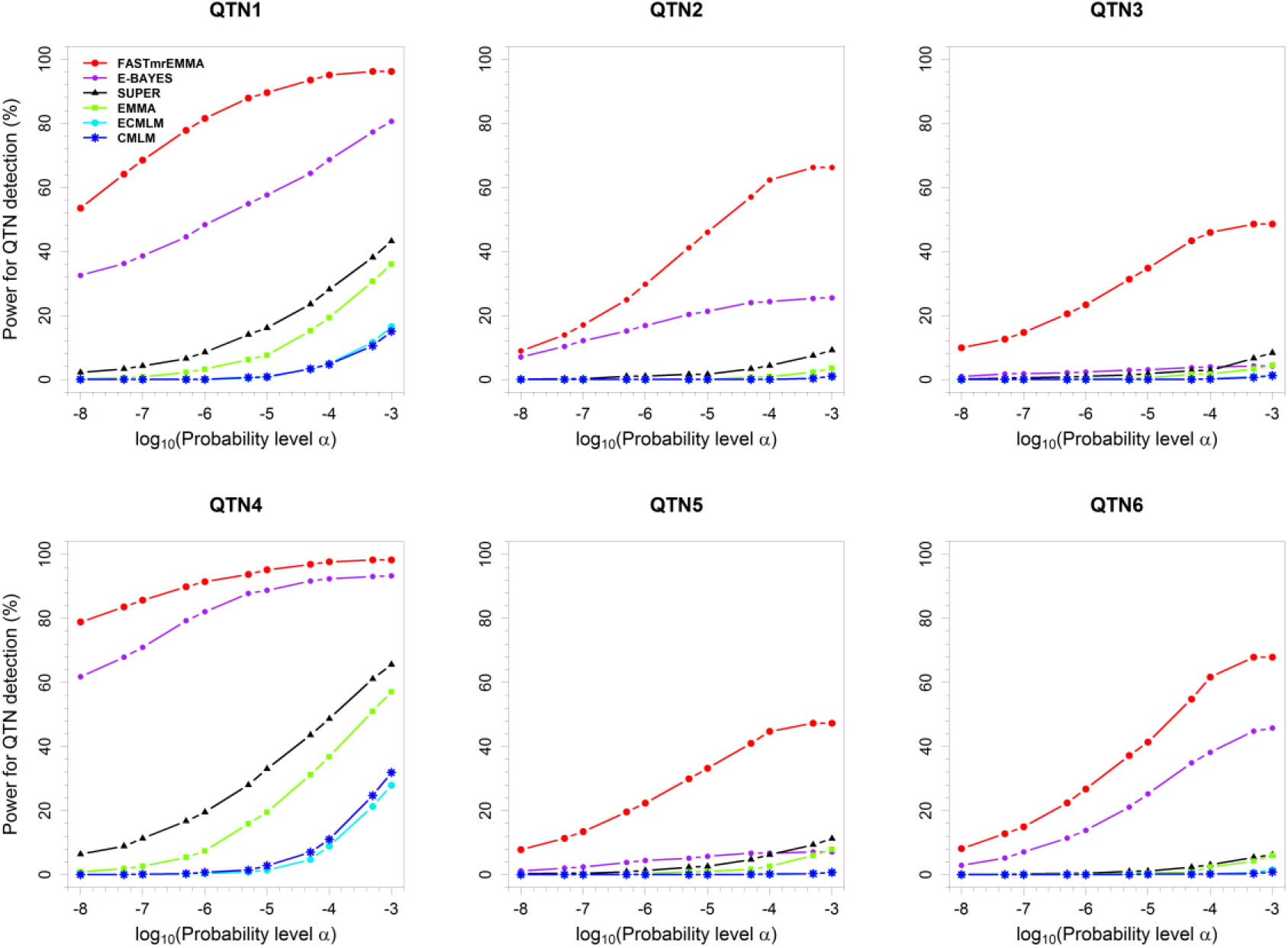
Statistical powers for six simulated QTN in the first simulation experiment plotted against type I error (in a log_10_ scale) for the six GWAS methods (FASTmrEMMA, E-BAYES, SUPER, EMMA, CMLM and ECMLM).

#### Computing time

In each of three simulation experiments, computing times for the six methods were recorded and are listed in **Table S4**. In summary, FASTmrEMMA has the least computing time followed by ECMLM, E-BAYES, CMLM and SUPER methods, and EMMA has the maximum computing time.

#### Real data analysis in Arabidopsis

To validate FASTmrEMMA, this new method along with E-BAYES, SUPER, EMMA, CMLM and ECMLM was used to re-analyze the *Arabidopsis* data [29] for days to flowering under long days (LD), days to flowering under short days with vernalization (SDV), leaf number at flowering with 8 wks vernalization, greenhouse (8W GH LN), and days to flowering, 8 wks vernalization, greenhouse (8W GH FT), and the results are listed in **Table S5**.

The numbers of SNPs significantly associated with the above four traits were 20, 17, 14 and 17, respectively, for traits LD, SDV, 8W GH LN and 8W GH FT, from FASTmrEMMA method. The corresponding numbers of the associated SNPs were 2, 6, 1 and 5 from E-BAYES; 21, 0, 0 and 0 from SUPER; 1, 5, 0 and 2 from EMMA; and 0, 1, 0 and 0 from both CMLM and ECMLM. Clearly, the number of significantly associated SNPs was much larger from FASTmrEMMA than from the other methods. These significantly associated SNPs for each trait were used to conduct a multiple linear regression analysis and the corresponding Bayesian information criteria (BIC) were calculated. For example, the BIC value for the model of 8W GH LN was −103.47 for FASTmrEMMA, 77.76 for E-BAYES and 117.50 for the others. FASTmrEMMA method shows the lowest BIC values for all the four traits (**Table 1**), indicating the best model fit among the six approaches.

**Table 1.** **Bayesian information criterion values for four flowering time traits in *Arabidopsis* using six genome-wide association study approaches**

| **Trait** | **FASTmrEMMA** | **E-BAYES** | **SUPER** | **EMMA** | **ECMLM** | **CMLM** |
| --- | --- | --- | --- | --- | --- | --- |
| LD | 39.54 | 287.00 | 396.65 | 299.97 | 382.07 | 382.07 |
| SDV | -88.09 | 43.20 | 179.54 | 100.69 | 169.87 | 169.87 |
| 8W GH LN | -103.47 | 77.76 | 117.50 | 117.50 | 117.50 | 117.50 |
| 8W GH FT | -321.72 | -155.55 | -82.41 | -101.83 | -82.41 | -82.41 |

LD: days to flowering under long days; SDV: days to flowering under short days with vernalization; 8W GH LN: leaf number at flowering with 8 wks vernalization, greenhouse;
8W GH FT: days to flowering, 8 wks vernalization, greenhouse.

Based on the SNPs detected by FASTmrEMMA, 6, 11, 5 and 7 genes were previously reported to be associated with the above four traits [32–34]. In the vicinity of the SNPs detected by E-BAYES, the corresponding numbers of the known genes are 2, 1, 0 and 1, respectively, for the above four traits [32]. Only four known genes for LD (SUPER), two known genes for LD (EMMA), and three known genes for SDV (EMMA) are in the neighborhood of the detected SNPs [32,34] (**Table 2**). Clearly, FASTmrEMMA method detected more known genes than did the other methods.

**Table 2.**
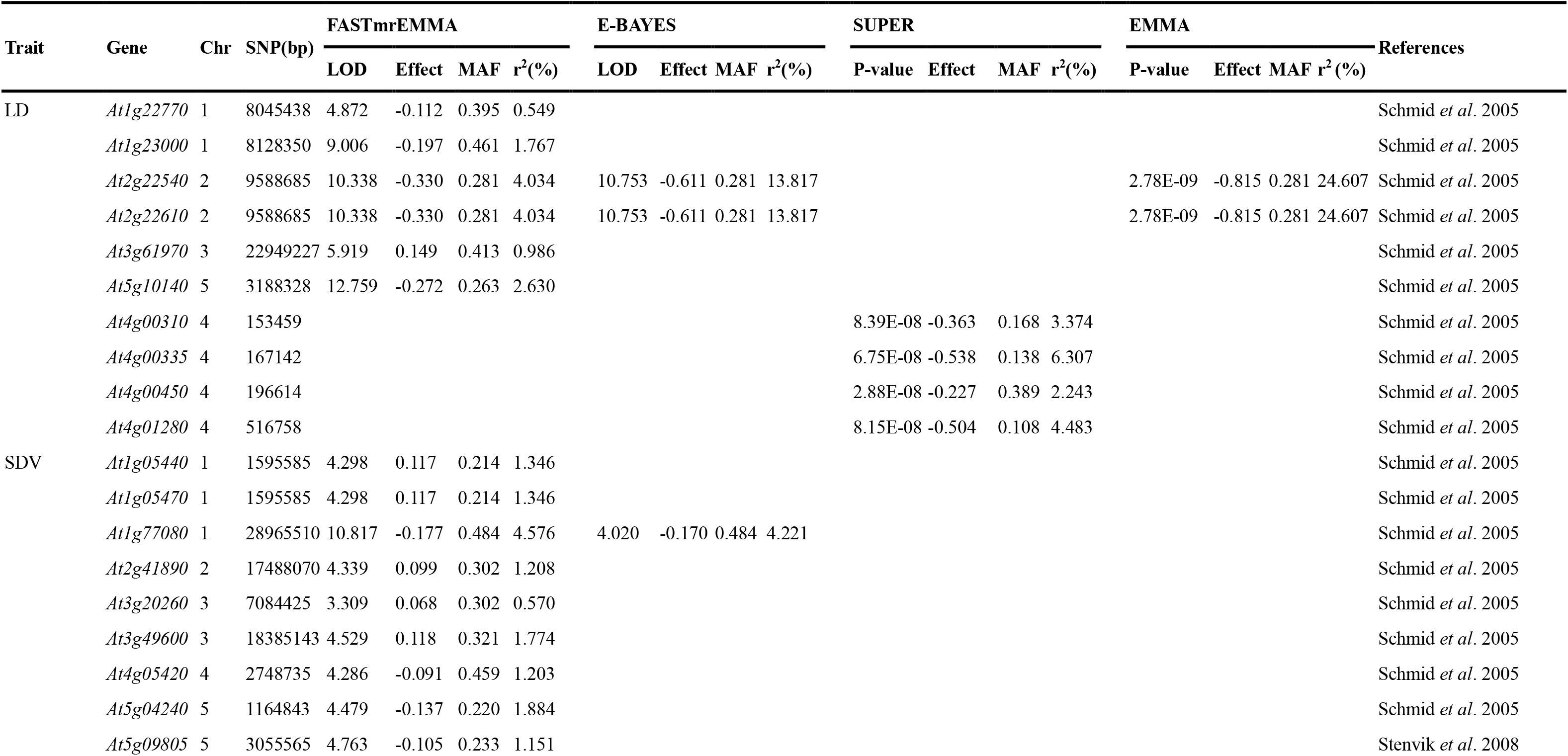
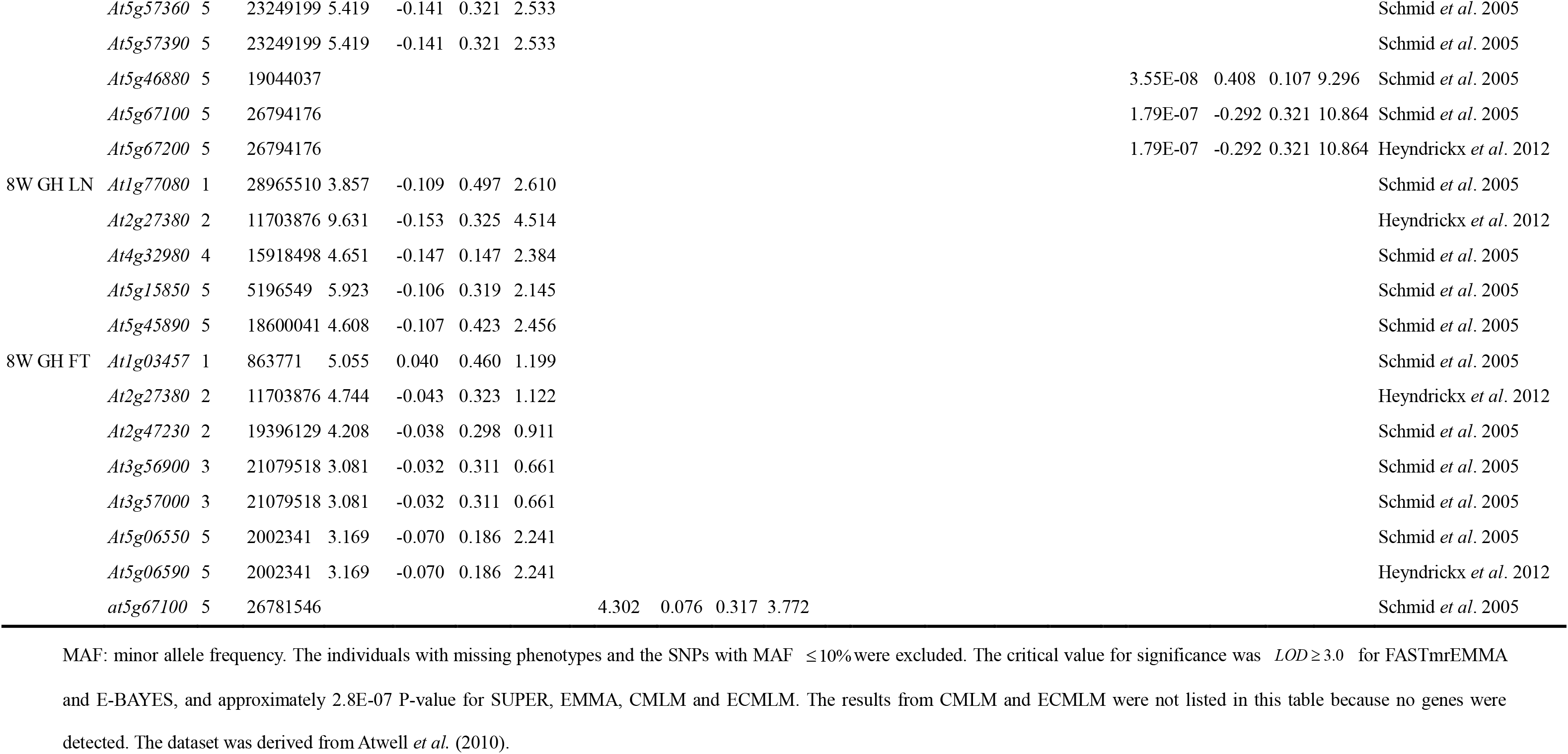
**Genome-wide association studies for four flowering time traits in *Arabidopsis* using six GWAS methods**

We also compared all the known genes detected in this study with all the candidate genes in Atwell *et al*. [29]. For example, among seven known genes (*At1g03457, At2g27380, At2g47230, At3g56900, At3g57000, At5g06550* and *At5g06590*) for 8W GH FT in this study, no genes were within the 133 candidate genes in Atwell *et al*. [29] (2010). Among 11 known genes for SDV in this study, only three genes (*At5g04240, At5g57360* and *At5g57390*) were within the 153 candidate genes in Atwell *et al.* [29].

## Discussion

When SNP effect is viewed as random, three variance components will be estimated. Generally, polygenic variance is larger than zero while variance components for most SNPs are zero, because these markers are not associated to the trait of interest. In other words, as in most mixed model approaches, variance components in FASTmrEMMA are also estimated under the assumption that one variance component is zero.

FASTmrEMMA is a new algorithm, and different from the widely-used one-dimensional genome scan approaches, such as SUPER, EMMA, CMLM and ECMLM. First, the SNP effects are viewed as random in FASTmrEMMA while they are viewed as fixed in SUPER, EMMA, CMLM and ECMLM. Because the random model approach will shrink the estimated SNP effects toward zero when the simulated QTN effects are small, leading to maximum correlation between observed and predicted phenotypic values [25,35]. Meanwhile, the power in the detection of QTN with random effect is higher than that with fixed effect [36].

Then, a quick single marker genome scan method was proposed to estimate the three variance components in the above mixed model. Here several techniques have been incorporated into the quick algorithm. The first technique is to fix the polygenic-to-residual variance ratio, which was adopted in CMLM/P3D [18] and EMMAX [20]. Although this algorithm is approximate, it has almost no effect on the estimate of SNP-effect variance, even if there is a large difference in the above ratios between the approximate and exact algorithms (**Table S1**). Clearly, this provides evidence for fixing the ratio in FASTmrEMMA. The second technique is to use a quick matrix calculation algorithm, such as, the eigen decomposition of matrix 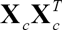 is the same as that of 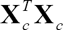 (a positive number). Thus, eigen decomposition, determinant and derivatives in the estimation of *λ*_*β*_ can be quickly calculated. The final technique is to estimate residual variance along with the estimation of fixed effects. In the single marker genome scan, therefore, only one parameter *λ*_*β*_ needs to be estimated so that running time is obviously decreased. Although GCTA algorithm [37] may be used to estimate the above three variance components, running time is a major concern. A similar situation is also apparent when using PROC MIXED in SAS in Zhang *et al.* (2005) [1].

Finally, our matrix transformation algorithm in FASTmrEMMA is different from those in SUPER, EMMA, CMLM, ECMLM and mrMLM^7^. For example, when many random effects are included simultaneously in one genetic model and polygenic background also needs to be controlled, at present there are no methods available. However, the new matrix transformation algorithm can transfer polygenic background plus residual error into a normal residual error. This new model can be easily treated by a Bayesian method. The applied study will be reported in the near future.

The multi-variance-component algorithm, E-BAYES [30], was also used to conduct multi-locus GWAS, especially for the situation where the number of markers is several times larger than sample size. However, results from simulation experiments showed that FASTmrEMMA is more powerful in QTN detection and higher accurate in QTN effect estimation than is E-BAYES. Note that E-BAYES method detected no genes in the real data analyses if all the 180,000 SNPs were simultaneously included in one genetic model (**Table S5**). To solve this issue, only a proportion of the SNPs, for example 2000 to 8000 SNPs, is included in one model at a time, so these SNP effects may be estimated by E-BAYES. FASTmrEMMA is different from the adaptive mixed LASSO [9]. If the number of markers is many larger than sample size, the adaptive mixed LASSO does not work. FASTmrEMMA is also different from both the Bayesian sparse linear mixed model [15] and the Bayesian mixture model [16]. The latter two operate under the framework of Bayesian statistics, and the computing time becomes a major concern.

FASTmrEMMA is different from MLMM of Segura *et al.* [17] in two aspects. First, MLMM is a simple, stepwise mixed-model regression with forward inclusion and backward elimination and FASTmrEMMA is a two-step combined method. In MLMM, the computationally-intensive forward-backward inclusion of SNPs is clearly a limiting factor in exploring the huge model space [17]. Second, matrix transformation algorithm in MLMM is different from that in FASTmrEMMA. This difference also exists between FASTmrEMMA and mrMLM of Wang *et al.* [7].

As described by Wang *et al.* [7], single-locus genome scan approaches for GWAS require Bonferroni correction for multiple tests. However, this correction often is too conservative to detect important loci for quantitative traits when the number of markers is extremely large. Clearly, FASTmrEMMA is based on a multi-locus model. Due to the multi-locus nature, Bonferroni correction is replaced by a less stringent selection criterion. Results from analysis of simulated and real data further validated the idea of a less stringent selection criterion in this study.

FASTmrEMMA is a combined method with two steps, each of which needs a critical P-value. In the first step, we compared the effect of three critical P-values (0.01, 0.005 and 0.001) on the power of QTN detection (**Table S6**) and found that the 0.005 critical P-value is the best. In the second step, a less stringent selection criterion between 0.05 and 0.05/*p* was adopted. The two critical P-values in FASTmrEMMA are confirmed by our simulated and real data analysis.

## Conclusion

In FASTmrEMMA algorithm, random-SNP-effect and multi-locus model methods are used to improve the power for QTN detection and to decrease the false positive rate, a new matrix transformation in the first step of FASTmrEMMA is constructed to obtain a new genetic model that includes only QTN variation and normal residual error. Additionally, letting the number of nonzero eigenvalues be one and fixing the polygenic-to-residual variance ratio are used to save running time. As a result, FASTmrEMMA has the highest power and accuracy for QTN detection and the best fit for a genetic model, as compared with E-BAYES, SUPER, EMMA, CMLM and ECMLM.

## Materials and Methods

### Genetic model

We consider the following standard MLM:

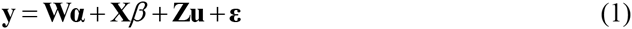

where **y** is an *n×*1 phenotypic vector of quantitative trait, and *n* is the number of individuals; 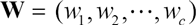 is an *n×c* matrix of covariates (fixed effects) including a column vector of 1, population structure [2] or principle component [39] may be incorporated into **W**, and **α** is a *c*×1 vector of fixed effects including the intercept; **X** is an *n*×1 vector of marker genotypes, and 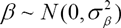 is random effect of putative QTN; **Z** is an *n*×*m* design matrix, 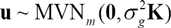 is an *m×*1 vector of polygenic effects; **K** is a known *m×m* relatedness matrix; and 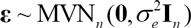 is an *n*×1 vector of residual errors, 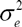 is the variance of residual error, **I**_*n*_ is an *n*×*n* identity matrix, and MVN denotes multivariate normal distribution. In animal datasets, *m* is the number of strains, *n* is the number of animals and **Z** indicates which strain each animal belongs to (*z*_*ij*_ = 1 if individual *i* comes from strain *j* and *z*_*ij*_ = 0 otherwise); and in the *Arabidopsis thaliana* data set of Atwell *et al.* [29], *m=n* and **Z = I**_*n*_.

In the current methods, including EMMA [3], CMLM/P3D [18], ECMLM [19], EMMAX [20], FaST-LMM [21], FaST-LMM-Select [22], SUPER [24], GEMMA [4], and GRAMMA-Gamma [23], *β* is treated as a fixed effect, from which it is relatively easy to estimate 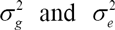. In this study we treat *β* as random in order to make the model more realistic [25,35,36]. In this case, three variance components need to be estimated under the assumption that QTN variance is zero, because most SNPs are not associated with the trait of interest. So the variance of **y** in the model (1) is

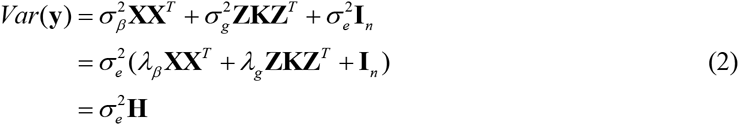

where 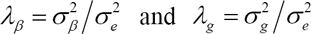.

### Fast multi-locus random-SNP-effect EMMA (FASTmrEMMA)

The key to solve the model (1) is to estimate 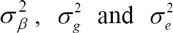. Although many algorithms or estimations are available, such as analysis of variance, maximum likelihood (ML), restricted maximum likelihood (REML), minimum norm quadratic unbiased, spectral decomposition [40] and average information [41], they are not feasible for a very high number of SNPs. Hence, we proposed a fast and efficient approximation algorithm in this study.

In the first step, we considered the reduced form of the model (1), which deleted **X***β*,

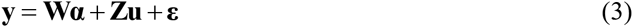

The variance of **y** is:

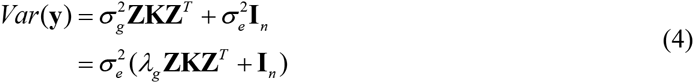

Using EMMA algorithm of Kang *et al.* [3], the estimate of *λ*_*g*_, denoted by 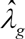, can be easily obtained.

In the second step, we considered the model (1), and replaced *λ*_*g*_ in (2) by the 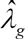, so

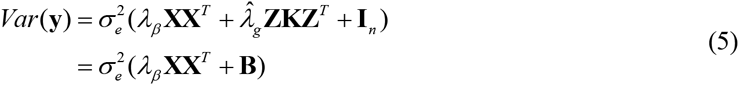

where 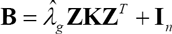. An eigen (or spectral) decomposition of the positive semi-definite matrix **B** was

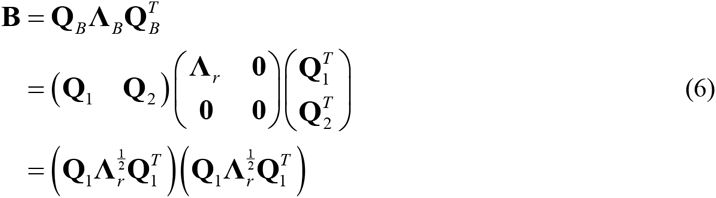

(**Appendix A**), where **Q**_*B*_ is orthogonal, **Λ**_*r*_ is a diagonal matrix with positive eigenvalues, *r* = *Rank*(**B**), **Q**_1_ and **Q**_2_ are the *n*×*r* and *n*×(*n−r*) block matrices of **Q**_*B*_, **0** is the corresponding block zero matrix.

Let 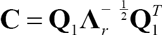, the model (1) was changed into

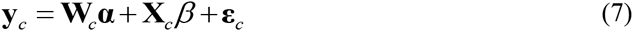

where 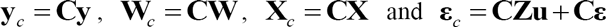. Clearly, the model (7) is a new 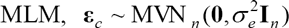, and

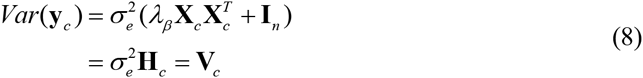

(**Appendix B**). Once the ratio of 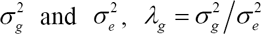, was fixed at 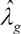, it is possible to scan each marker on the genome. The evidence for the effectiveness of this approximation is shown in the results section.

#### Log-likelihood and restricted log-likelihood functions

According to the descriptions for the single-locus genome scan algorithm in previous GWAS studies [3,4,43,44], log-likelihood and restricted log-likelihood functions for the model (7) are

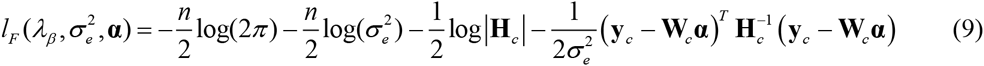

and

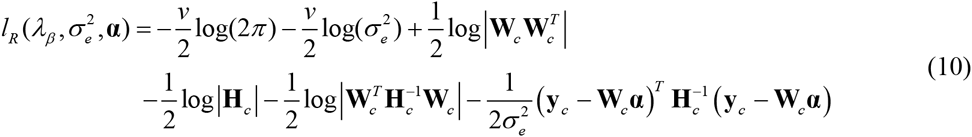

respectively, where 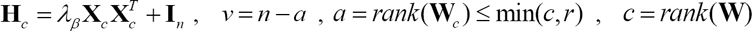 *r = rank*(**C**) = *rank*(**B**), supposing **W** is column full rank.

Once **α** and 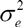 are fixed, the ML and REML estimates for *λ*_*β*_ is equivalent to maximizing the following target functions

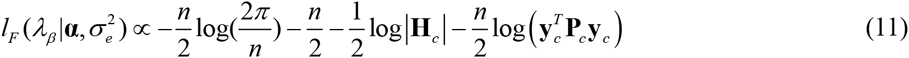

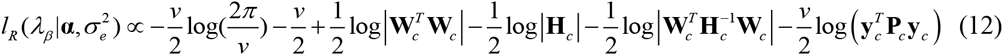

where 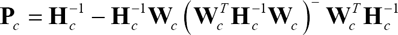, and – denotes generalized inverse.

Since it is very slow to calculate determinant and inversion in the equations (11) and (12), a fast computation algorithm should be considered. As described in GEMMA, Zhou and Stephens [4] first obtained the first and second derivatives for *λ*_*β*_, and then conducted eigen (or spectral) decomposition. In EMMA, however, Kang *et al.* [3] first conducted eigen decomposition, and then calculated the derivatives. These two ways are essentially the same. For simplicity, we adopted EMMA method.

It is possible to find *ξ*_*i*_ and *δ*_*s*_, such that

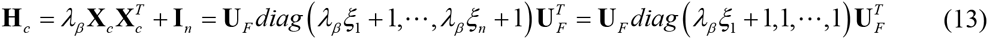

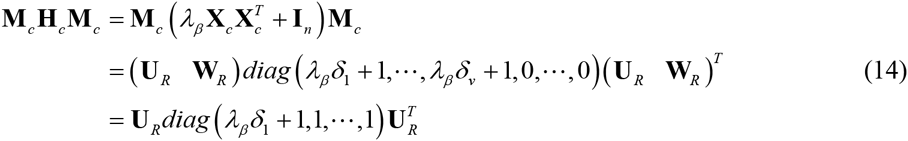

where 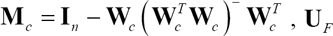 is an *n*×*n* orthogonal matrix, and **U**_*R*_ is an *n×v* eigenvector matrix corresponding to the nonzero eigenvalues, and **W**_*R*_ is an *n×*(*n−v*) eigenvector matrix corresponding to zero eigenvalues.

Note that *ξ*_*i*_ = 0 (*i* = 2,…,*n*) and *δ*_*s*_ = 0 (*s* = 2,…,*v*). This is because the nonzero eigenvalues of 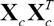 are the same as those of 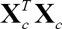, which is a positive number. For technical detail the reader is referred to **Appendix C**. It should be noted that ***U***_*F*_ and **U**_*R*_ are independent of *λ*_*β*_. Let 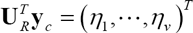, then finding the ML and REML estimates for *λ*_*β*_ is equivalent to optimizing the following functions with respect to *λ*_*β*_ (**Appendix D**):

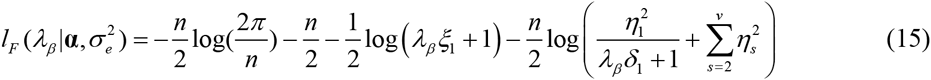

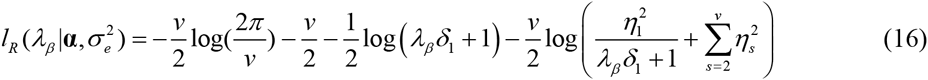

#### Estimation of parameter *λ*_*β*_

At present three algorithms, Newton-Raphson (NR), Fisher scoring and expectation-maximization, are frequently used to obtain the ML and REML estimates [45]. In this study we adopted the NR algorithm, which has the form

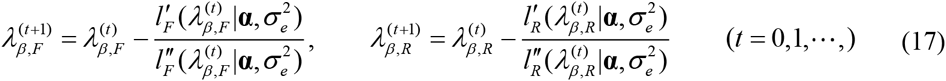

where 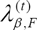 and 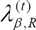 are the ML and REML estimates at the *t*th iteration, respectively; and 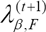 and 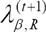 are the update estimates, respectively, and the first derivatives of these two functions on *λ*_*β*_ were

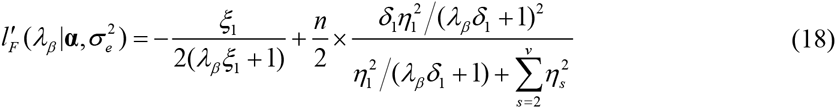

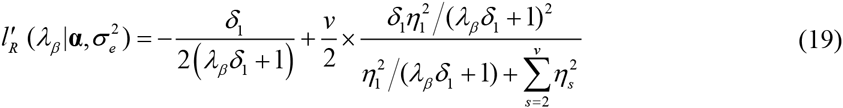

and the second derivatives on *λ*_*β*_ were

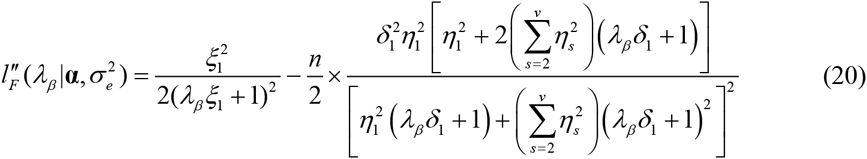

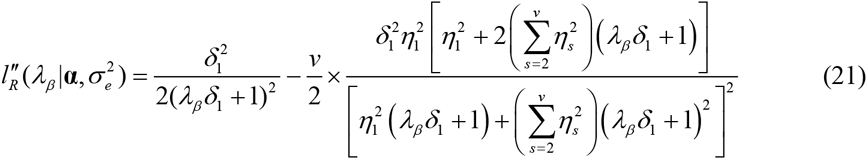

Using the idea of EMMA [3], the range of *λ*_*β*_ is between 1E-05 (corresponding to almost pure environmental effect) to 1E+05 (corresponding to almost pure single-gene effect), and we divided this range evenly into 100 regions in logarithm scale to compute (18) or (19). The global ML or REML is searched for by applying the NR algorithm to all the intervals where the signs of derivatives change. This optimization technique for estimating *λ*_*g*_ has guaranteed the convergence as long as the kinship matrix **K** is positive semi-definite. Note that 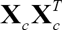 is always positive semi-definite.

#### Estimation of fixed effects *α* and residual variance 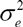

Once *λ*_*β*_ is known, it is easy to estimate **α** and 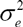. The ML estimates for **α** and 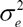 were

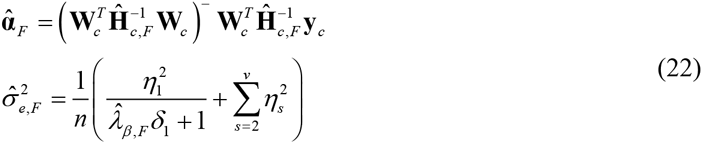

respectively, where 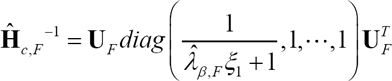. The REML estimates for **α** and 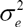 were

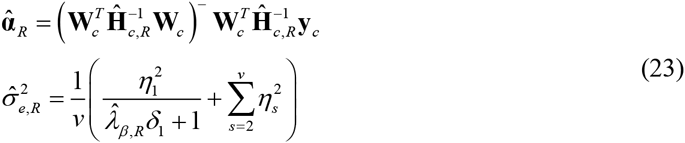

respectively, where *v* = *n* –*a* and 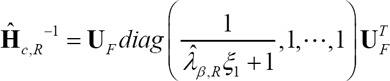 (**Appendix D**).

#### Best linear unbiased prediction (BLUP) for parameter *β*

Using 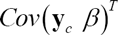, the BLUP for the 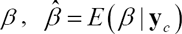, can be obtained. Based on the above fast and efficient algorithm, the ML or REML estimate is

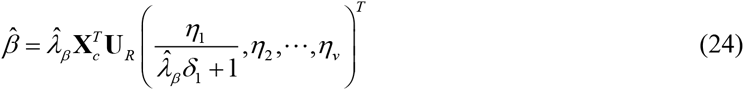

(**Appendix E**), where 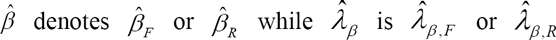, respectively.

#### Likelihood ratio test (LRT)

Although the parameter on QTN in the model (1) is 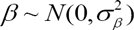, the estimation of *λ*_*β*_ is our concern in the above algorithm. Therefore, the null hypothesis might be *λ*_*β*_ = 0 [35]. The LRT statistic for the ML or REML estimate is

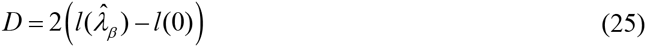

where 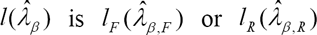, and 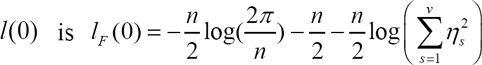, or 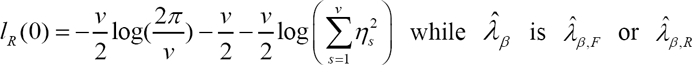, or respectively.

Under the null hypothesis, the LRT statistic *D* follows approximately a mixture of two 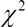 distributions with an equal weight, denoted by 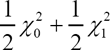, where 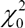 is just a fixed number of zero and 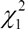 is a 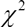 distribution with one degree of freedom [35], and the P-values can be calculated accordingly. Let P be the P-value for each QTN, it was calculated using

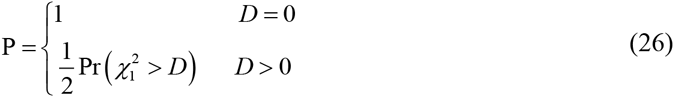

#### Kinship matrix

Many methods for calculating kinship matrix **K**_*m×m*_ from a large number of markers have been proposed, such as identical-by-state approach [2,3,7,26,46]. Here we adopted the method of Kang *et al.* [3]. Let *S* be an *p*×*m* genotypic matrix with elements 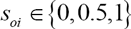, the element of kinship matrix **K** is defined by

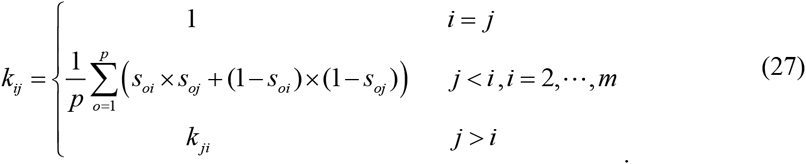

#### Time complexity for the first step of FASTmrEMMA

For the single-locus genome scan, FASTmrEMMA is involved in only one eigen decomposition at the beginning, and computational complexity is *O*(*mn*^2^), where *O* is the big *O* notation. For each SNP tested, FASTmrEMMA effectively replaces the expensive additional eigen decomposition step in EMMA, and the computational complexity changes from 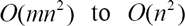, because the nonzero eigenvalues of 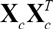 are the same as those of 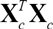. After this, as in EMMA, each iteration of the optimization step requires inexpensive operations (computational complexity of *O*(*n*)) to evaluate both the first and second derivatives of target functions. Therefore, the overall time complexity for the first step of FASTmrEMMA is *O*(*mn*^2^ + *pn*^2^ + *ptn*), compared with *O*(*mn*^2^ + *pmn*^2^ + *ptn*) for EMMA[4], where *t* is the number of optimization iterations required for the Newton-Raphson method (quadratic rate of convergence).

In the GWAS, the number of SNPs is often 1000 times larger than the sample size, and actually most SNPs are not associated with the trait of interest. In this case, fitting all the genome markers in one model is not feasible. Once we delete these SNPs with zero effects, the reduced model is estimable. The described above can be considered as an initial screening step for FASTmrEMMA. In the first step of FASTmrEMMA, a less stringent criterion for the initial stage screening was adopted, for example, all the SNPs with the ≤0.005 P-values were selected to enter the next step for further evaluation. The majority of markers will be eliminated in the first step. Therefore, the number of markers left in the second stage analysis is often a small subset of all markers, say a few hundred or a few thousand at most, for example, no more than 600 significantly associated SNPs for the four traits in the *A. thaliana* datasets [29].

In the multi-locus model, we proposed to use the EM empirical Bayes (EMEB) [28] because EMEB method is a random model approach in which each random marker effect is assigned an empirical distribution with a variance, and therefore is in accordance with treating a marker as a random effect in this study. The linear model is as followed:

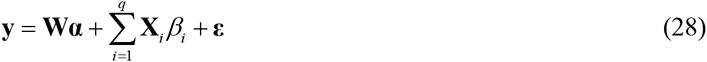

where **y**,**W**, **α** and **ε** are the same as model (1); *q* is the number of the selected QTN in the first step of FASTmrEMMA; **X**_*i*_ and *β*_*i*_ are an *n×*1 vector of marker genotypes and effect for the *i* th QTN, respectively. In the above model, polygenic background is not included, because all the potential QTN have been included in the model (28).

In the model (28), we adopt the normal prior for 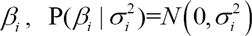, and the scaled inverse 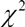 prior for 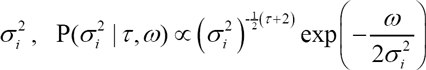, where we set (*τ*,*ω*) = (0,0), which represents the Jeffreys’ prior, 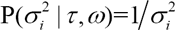 [47]. The procedure for parameter estimation in EMEB [28] is as follows.

1) Initial-step: To initialize parameters with

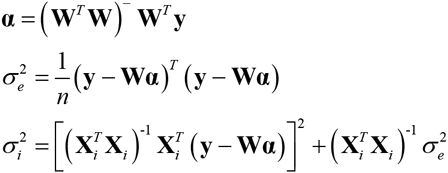

2) E-step: QTN effect can be predicted by

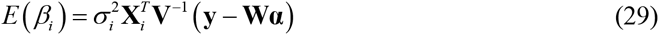

where 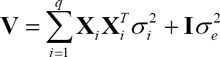.

3) M-step: To update parameters 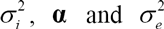:

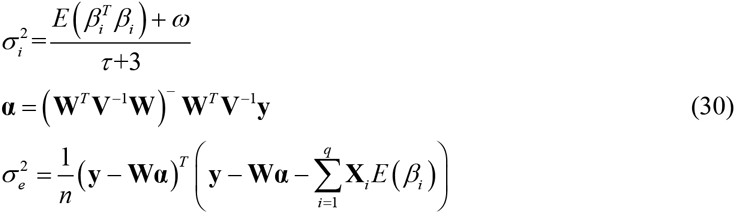

where 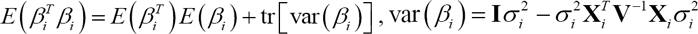 and 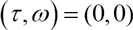. Repeat E-step and M-step until convergence is satisfied.

Because the model is multi-locus in nature, Bonferroni correction is replaced by a less stringent selection criterion. Although the general 0.05 critical value may be used for the significance test, we decided to place a slightly more stringent criterion of LOD=3.0. The criterion is frequently adopted in linkage analysis and is the equivalent of 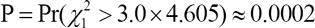, in which 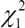 under the null hypothesis, follows a 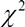 distribution with one degree of freedom.

### Other five methods

E-BAYES (see the Additional Information) is an existing multi-locus Bayesian approach implemented by the SAS program[30], and was used as a gold standard for multi-locus model comparison.

EMMA, CMLM, ECMLM and SUPER (see the Additional Information) were used as gold standards for single-locus model comparisons. All the thresholds of P-values for significance tests were set at 0.05/*p*, due to Bonferroni correction for multiple tests, where *p* is the number of markers.

### The Arabidopsis thaliana data

We analyzed the well-known *A. thaliana* datasets published by Atwell *et al.* [29]. Both phenotypes and genotypes were obtained from http://www.arabidopsis.usc.edu/. A total of 199 *Arabidopsis* lines and 216130 SNPs were used for analysis. Four flowering time traits (LD, SDV, 8W GH LN and 8W GH FT) with log-transformation were re-analyzed in this study. We excluded the individuals with missing phenotypes, non-polymorphic SNPs and SNPs with minor allele frequency (MAF) less than 10% and all the six methods (FASTmrEMMA, E-BAYES, SUPER, EMMA, CMLM and ECMLM) were used to analyze these four datasets. A total of approximate 180,000 SNPs for each trait were used to calculate the identity by state (IBS) matrix as the estimates of relatedness [3].

### Simulation experiments

Three Monte Carlo simulation experiments were conducted to validate the new algorithms.

As described by Wang *et al.* [7] (2016), the SNP genotypes derived from the *A. thaliana* datasets [29] were also used to perform three simulation experiments. The purpose was to compare FASTmrEMMA with the single-locus model methods (SUPER, ECMLM, CMLM and EMMA) and the multi-locus model method (E-BAYES). In the first simulation experiment, 2000 SNPs on each chromosome were randomly sampled. As a result, all the SNPs between 11226256 and 12038776 bp on Chr. 1, between 5045828 and 6412875 bp on Chr. 2, between 1916588 and 3196442 bp on Chr. 3, between 2232796 and 3143893 bp on Chr. 4, and between 19999868 and 21039406 bp on Chr. 5 were used to conduct simulation studies. The sample size was 199, the number of lines from Atwell *et al.* [29]. Six QTNs were simulated and placed on the SNPs with allele frequencies of 0.30; their heritabilities of each effect size were set as 0.10, 0.05, 0.05, 0.15, 0.05 and 0.05, respectively; and their positions and effects are listed in Table S2. The residual variance was set at 10.0. The new phenotypes were simulated by the model: 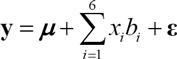, where 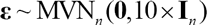. Each sample was analyzed by the above six methods. For each simulated QTN, we counted the samples in which the LOD statistic exceeded 3.0 for FASTmrEMMA, the P-value was less than 0.05 for E-BAYES, and the P-value ≤5e-6 (0.05/*p*) for the others. A detected QTN within 2 kb of the simulated QTN was considered a true QTN. The ratio of the number of such samples to the total number of replicates (1000) represented the empirical power of this QTN. The Type I error was calculated as the ratio of the number of false positive effects to the total number of zero effects considered in the full model. To measure the bias of QTN effect estimate, mean squared error (MSE) and mean absolute deviation (MAD) were defined as

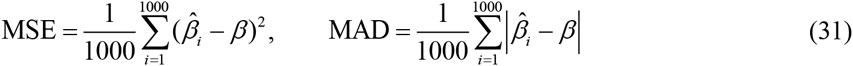

where 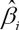 is the estimate of *β* for each QTN in the *i*th sample. A method with a small MSE (or MAD) is generally more preferable than a method with a large MSE (or MAD).

To investigate the effect of polygenic background on FASTmrEMMA, polygenic effect was simulated in the second simulation experiment by multivariate normal distribution 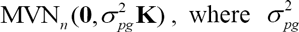 is polygenic variance, and **K** is the kinship coefficient matrix between a pair of lines. Here 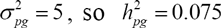. The QTN size (r^2^), residual variance, and others were the same as those in the second simulation experiment. The new phenotypes were simulated by the model: 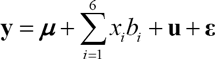, where polygenic effect 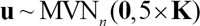 and 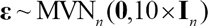. To investigate the effect of epistatic background on FASTmrEMMA, three epistatic QTN each with 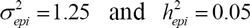 were simulated in the third simulation experiment. The first one was placed between 3063784 bp on Chr. 4 and 5227063 bp on Chr. 2; the second one was placed between 5986135 bp on Chr. 2 and 2031781 bp on Chr. 3; and the third one was placed between 2668059 bp on Chr. 3 and 11824678 bp on Chr. 1. The QTN size (r^2^), residual variance and others were also the same as those in the first simulation experiment. The new phenotypes were simulated by the model: 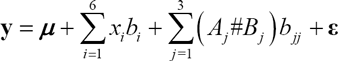, where 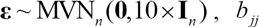 is epistatic effect, and 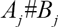 is its incidence coefficient.

## Acknowledgements

This work was supported by the National Natural Science Foundation of China (grants 31571268 and 31301229), and Huazhong Agricultural University Scientific & Technological Self-innovation Foundation (Program No. 2014RC020).

## Authors’ contributions

Y.-M.Z. conceived and supervised the study, and revised the manuscript. R.W. assisted the supervision of the experiments and revised the manuscript. Y.-J.W. deduced the formulas and wrote the draft. Y.-J.W., S.-B.W., H.Z., B.H., J.Z. and J.-Y.F. performed the experiments and analyzed the data. J.M.D. and H.Z. improved the language within the manuscript. All authors read, revised and approved the final manuscript.

## Additional Information

### Competing Financial Interests

The authors declare no competing financial interests.

### Supplementary information

accompanies this manuscript in the file entitled with “Additional Information”.

